# Three linked opposing regulatory variants under selection associate with *IVD*

**DOI:** 10.1101/2022.12.22.521605

**Authors:** Elizabeth A. Brown, Susan Kales, Michael J. Boyle, Joseph Vitti, Dylan Kotliar, Stephen F. Schaffner, Ryan S. Tewhey, Pardis C. Sabeti

**Author notes:** These individuals co-supervised the work (Ryan S. Tewhey, Pardis C. Sabeti). This is the corresponding author.

## Abstract

While genome-wide association studies (GWAS) and selection scans identify genomic loci driving human phenotypic diversity, functional validation is required to discover the variant(s) responsible. We dissected the *IVD* locus, implicated by selection statistics, multiple GWAS, and clinical genetics as important to function and fitness. We combined luciferase assays, CRISPR/Cas9 genome-editing, massively parallel reporter assays (MPRA), and bashing of regulatory loci. We identified three regulatory variants, including an indel, that may underpin GWAS signals for pulmonary fibrosis and testosterone, and that are linked on a positively selected haplotype in the Japanese population. These regulatory variants exhibit synergistic and opposing effects on *IVD* expression experimentally. Alleles at these variants lie on a haplotype tagged by the variant most strongly associated with *IVD* expression and metabolites, but with no functional evidence itself. This work demonstrates how comprehensive functional investigation and multiple technologies are needed to discover the true genetic drivers of phenotypic diversity.

## INTRODUCTION

Whole genome^1^ and exome^2^ sequencing of thousands of individuals has yielded rich maps of human genetic variation, data that should in principle permit systematic study of how genotype affects phenotype. In practice, however, linking these two is difficult, and research still focuses largely on the variation that is easiest to study: single nucleotide polymorphisms (SNPs) that affect protein structure and that have large impacts on human disease^2,3^. These protein-coding SNPs only explain a small fraction of heritable variation in phenotypes^4^. New approaches are needed to tackle the contribution of complex human genetic variation to phenotype, as it plays out for the majority of associated traits.

To fully elucidate the loci that govern critical traits, we will need to probe genetic variation more broadly, including (1) non-coding regulatory variation, (2) different classes of mutation beyond SNPs, and (3) combinations of multiple variants within a single locus. First, it is critical to study non-coding regulatory variation because GWAS and genomic scans for natural selection implicate theseregions as the predominant drivers of biological diversity^5,6^. Second, in depth analyses of regulatory regions must query insertions and deletions (indels), repeats, and translocations in addition to SNPs, which have been the focus purely due to their simplicity in sequencing and association studies. Third, multiple variants may modulate activity of a gene, controlling a single phenotype in concert^4,7^; they must therefore be studied together in varied combinations. For example, a study of autoimmune GWAS loci found patterns consistent with multiple linked variants controlling gene expression at these loci^8^.

To understand complex loci we implemented a process to move from genomics to function at critical regulatory loci (Supplementary Fig. 1), specifically to dissect *cis* regulation of *IVD* (isovaleryl-CoA dehydrogenase) after identifying it as a candidate for recent positive selection. *IVD* acts in catabolism of leucine, an essential amino acid in our diet with roles in muscle anabolism, fatty acid oxidation, and insulin signaling early in development^9–12^. The locus came to our attention when a genome-wide scan for positive selection using the Composite of Multiple Signals (CMS), which combines several population genetic statistics of selection, identified a significant signal around *IVD* in the Japanese (JPT) population from phase 3 of the 1000 Genomes Project (1000G)^6,13^.

Several lines of evidence, beyond the signal of positive selection, suggested that variants at *IVD* may have important impacts on fitness. First, there is evidence that genetic variants regulate *IVD* expression, and also cellular metabolite levels. Variation in the region strongly associates with *IVD* expression, and also with levels of the leucine metabolite and *IVD* substrate, isovalerylcarnitine in metabolomic GWAS and RNA-seq studies^14,15^. Second, there is evidence that non-coding variation in *IVD* impacts health. Non-coding variants of *IVD* are associated with pulmonary fibrosis^16,17^ and lung function - vital capacity and forced expiratory volume - in a large GWAS of Europeans^18^. Non-coding variants in the region also associate with measures of intelligence^19–24^ and testosterone^25,26^ across a number of GWAS in Europeans. *IVD* coding variants, on the other hand, cause isovalerylcarnitine to accumulate in a rare disease called isovaleric acidemia^9,27^, and mouse knockouts are homozygous lethal^28^. Without dietary restriction of leucine, isovaleric acidemia can manifest as acute/lethal neonatal seizures and vomiting, chronic intermittent disease, or mild forms depending on the mutation that may result in less severe metabolic symptoms and cognitive impairment^29^.

We used multiple statistical and experimental approaches to detect, validate, and characterize *IVD* eQTLs regulating leucine metabolism. Specifically, we started by testing whether variants in the association peaks altered gene expression, using traditional luciferase reporter assays^30^. We then used CRISPR/Cas9-based homology directed repair (HDR)^31,32^ to replace alleles and verify endogenous regulation of *IVD*. As high-throughput technologies became available, we used a multiple parallel reporter assay (MPRA)^33–35^ to test associated variants across the *IVD* region more broadly, and followed up additional functional candidates with luciferase assays. We further attempted to localize regulatory activity at individual nucleotides by using an MPRA bashing strategy of tiling deletions across a regulatory region. Alongside our functional tests, we queried the peaks of association more deeply, searching for tightly-linked, but independent causal regulatory variants. Lastly, we explored signals of natural selection and emergence of *IVD*’s regulatory variants and haplotypes. Our work lays the foundation for understanding how multiple regulatory variants govern a selected region associated with pulmonary function, and subject to monogenic disease in humans.

## RESULTS

### Analyzing IVD eQTLs, isovalerylcarnitine association, and selection peak

Our first step was to examine whether regulatory variation in the *IVD* region could potentially impact phenotype or fitness. Since variants in the *IVD* locus strongly associate both with *IVD* expression and levels of its substrate, isovalerylcarnitine^14,15^, we tested whether the variants most strongly associated with *IVD* expression correlate with those associated with isovalerylcarnitine levels. We found positive correlation between these association peaks (r^2^ = 0.40, p-value < 2e-16 for the slope), suggesting *cis*-regulation of *IVD* does impact its catalytic activity and levels of its metabolite (Fig. 1 and Supplementary Tables 1 and 2).

**Figure 1.**
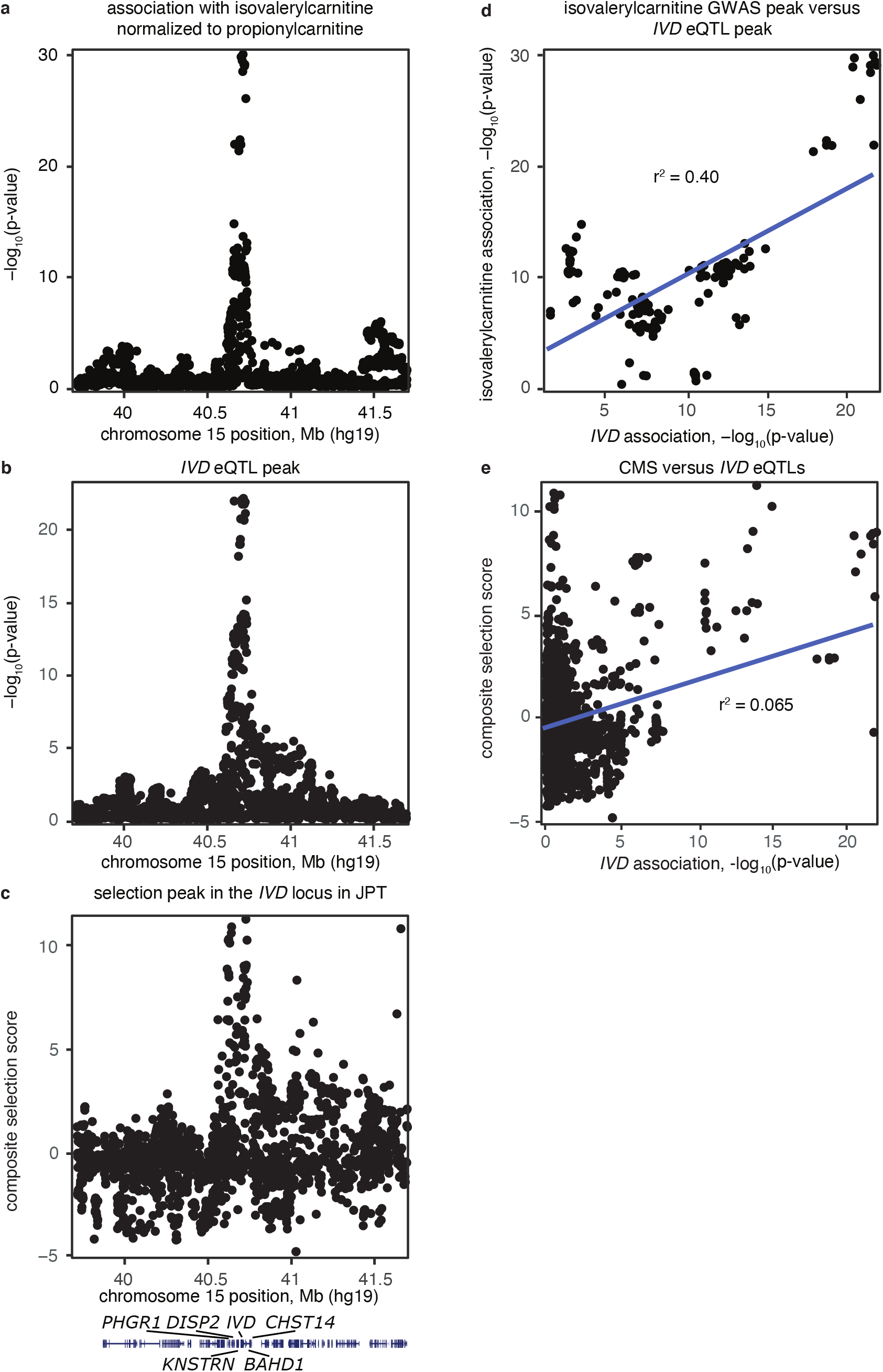
*IVD* variants associate with *IVD* expression, isovalerylcarnitine, and selection in JPT. **a** Variation in the *IVD* locus strongly associates with isovalerylcarnitine, the metabolite in the leucine catabolism pathway, which the IVD enzyme catalyzes to 3-methylcrotonylcarnitine. Metabolomics and genotyping was conducted in 4,600 individuals from the Twins UK cohort, and isovalerylcarnitine levels are normalized to propionylcarnitine. **b** Variation in the *IVD* locus strongly associates with IVD expression in LCLs from 373 European individuals in the Geuvadis dataset. **c** A selection peak centers around the *IVD* locus in the JPT population. The variant scores are based on a normalized composite of multiple signals (CMS). **d** Association of variants with isovalerylcarnitine (normalized to propionylcarnitine) correlated significantly with their association with IVD expression in LCLs from the Geuvadis study. From linear regression analysis y = 0.77109*x – 5.74396 with p < 2e-16 and r^2^ = 0.40. **e** CMS scores in the *IVD* region in the JPT population correlated weakly but significantly with their association with *IVD* expression in LCLs from the Geuvadis study. From linear regression analysis y = 0.22828*x - 0.30910 with p < 2e-16 and r^2^ = 0.065.

Given the signal of recent positive selection in the JPT population at this locus, we also examined whether regulatory variation in the region might have fitness implications. To do this we compared the CMS score in JPT to the association with IVD expression (detected in a European population) at each variant. We found positive correlation between signatures of selection and association with *IVD* expression (r^2^ = 0.065, p-value < 2e-16 for the slope) (Fig. 1 and Supplementary Table 3). These positive correlations between *IVD* eQTLs, isovalerylcarnitine metabolite levels, and signals of recent positive selection in JPT gave us reason to believe that regulation of *IVD* is important to phenotype and fitness, and therefore worthy of functional dissection to identify and validate active regulatory variation.

### Luciferase reporter assays

To identify potential regulatory variants in the *IVD* locus, we designed luciferase assays to probe a small tractable set of most likely candidates within a large genomic context. Looking closely at the region surrounding the top *IVD* function candidates, we observed tracks of DNase I hypersensitivity and histone modifications that extended hundreds of basepairs. We recognized that for our assays to be most successful, we would want to include large regions to maintain relevant genomic context for enhancer function. Given the manual nature of such an effort, we selected a set of eight SNPs linked with r^2^ > 0.9 to the peak associated *IVD* eQTL (rs11638033, referred to here as “τ”) and GWAS SNP.

We used DNase I hypersensitivity and histone modifications, including H3K4Me1 often found near enhancer elements, as guides to select seven regions each of 600-bp – 2-kb in size encompassing these eight candidate variants for examination (Fig. 2a). Six of the regions each harbored 1 of the variants, and one harbored two of the variants, rs17733719 and rs8033249. We cloned out ancestral and derived haplotypes for each of these regions from a lymphocyte cell line (LCL), NA12812, which is notably heterozygous for all eight candidates. In the case of rs17733719 and rs8033249, we used site directed mutagenesis to convert the ancestral variants to derived, individually, in order to test them separately. We found that just one of the seven tested regions, the 2kb region falling 3’ of *IVD* coding region, showed a phenotypic effect. The region, which contained a proxy for the τ variant (rs11633883) and nine additional heterozygous sites, had significantly increased expression for the derived haplotype compared to the ancestral haplotype (Fig. 2b).

**Figure 2.**
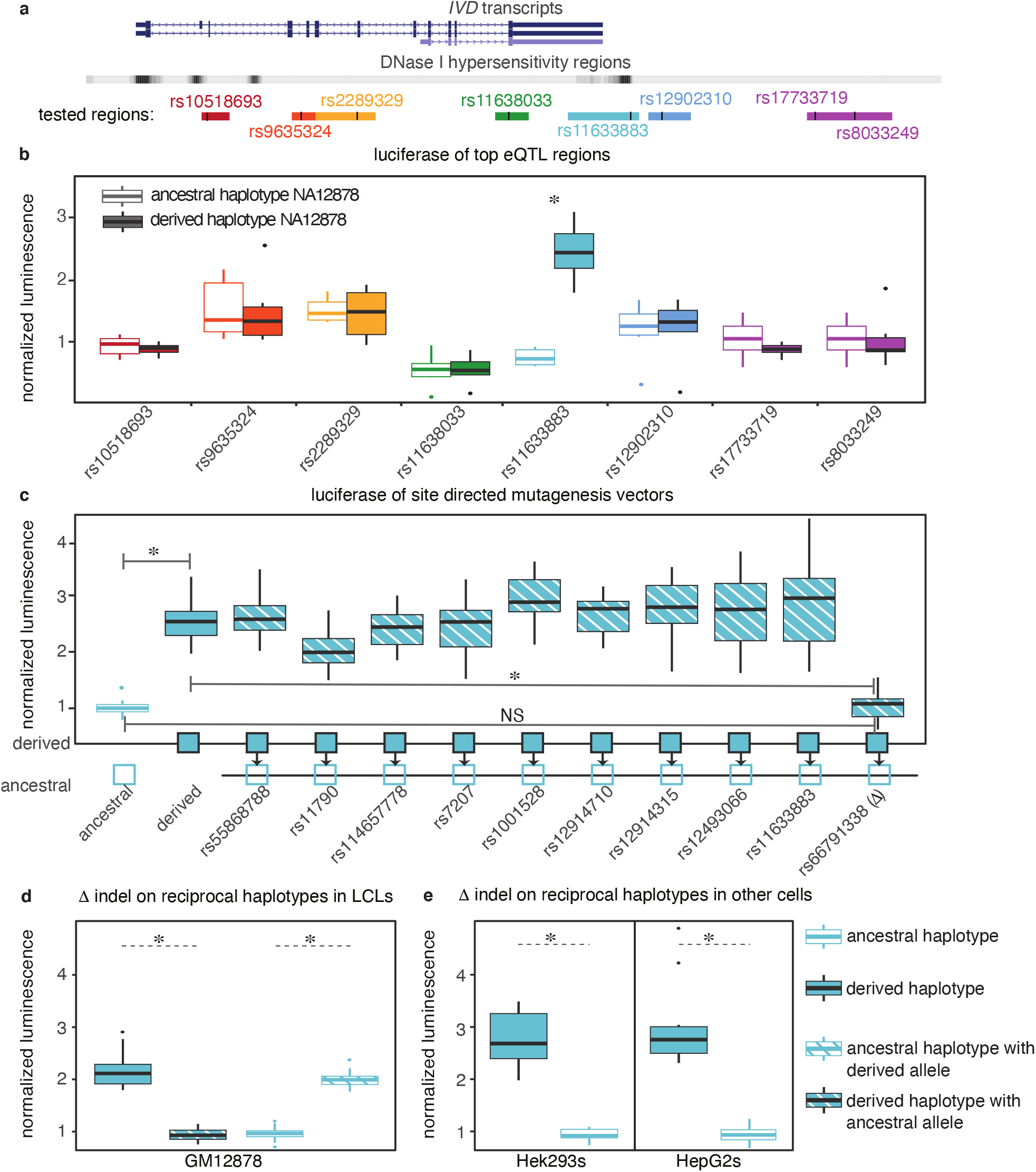
Luciferase assays identify a 5-bp indel with regulatory capacity. **a** We created pGL4.23 constructs that included 600bp - 2kb long regions surrounding each of our 8 variants of interest, cloned from NA12878 LCLs. We denote each region by color and show it with respect to the IVD transcript and DNaseI hypersensitivity sites in the region. We mark variants of interest with thick black lines in each region. rs17733719 and rs8033249 fall in one region, so the same ancestral haplotype data is the same for each, and we tested derived alleles of these separately after using site-directed mutagenesis. **b** We plot luminescence of the pGL4.23 firefly luciferase for each construct using boxplots, normalized to the pGL4.74 renilla luciferase spike-in (normalized luminescence). Constructs containing derived variants are solid boxes, and ancestral are opaque. **c** We used site directed mutagenesis to convert variants from derived to ancestral on this 2kb haplotype. We show normalized luminescence of each of these 10 mutated constructs in comparison to the ancestral and derived constructs. **d** We reciprocally convert Δ from derived to ancestral and ancestral to derived on the derived and ancestral haplotype backgrounds in pGL4.23 vector backbone, and test how this recapitulates the change in activity level by luciferase assay in GM12878 LCLs. **e** We also test the impact of the ancestral and derived haplotypes in HepG2s and Hek239s. *indicates p-value < 0.01 for 2-sample t-test.

Having identified one 2kb region that produced differences in expression between the two tested haplotypes, we next sought to discriminate which of the ten linked loci in the region was responsible for the change. To accomplish this, we began with the 2kb cloned out region in NA12812 LCLs and converted all ten variants, one at a time, on the derived haplotype to the alternative allele on the ancestral haplotype. We found that just one of the ten variants had an observed phenotypic effect. The variant, a 5-bp indel (rs66791338, referred to here as “Δ”), reduced luciferase expression when it was changed from derived deletion (-) to ancestral insertion (GAAAG) (Fig. 2c). The Δ indel has r^2^ = 0.6 in GBR to τ, the top associated eQTL in the European population. None of the other nine variants, including the more strongly linked variant (r^2^ = 0.98 in GBR), had any effect when perturbed.

We further focused on confirming the effect of Δ variant in modulating reporter expression on a vector backbone, and testing its effect in other cell lines. Reciprocally, converting Δ from ancestral to derived recapitulates the change in activity level by luciferase (Fig. 2d). The change in activity when testing the Δ indel alone mirrors the observed change for the full haplotype. Furthermore, this change in activity was consistent across two other cell types we tested, hepatocytes (HepG2) and embryonic kidney cells (HEK293) (Fig. 2e).

### CRISPR/Cas9 endogenous replacement of Δ indel

Next, we tested whether the 5-bp indel (Δ) impacts expression of *IVD* endogenously in LCLs using CRISPR/Cas9 HDR to convert Δ from homozygous ancestral to homozygous derived and vice-versa. We isolated and identified individual clones either homozygous for HDR of the Δ indel, or homozygous for no change of the indel. After growing the cells from clones, we measured *IVD* expression relative to two housekeeping genes (PPIA and HSPCB). We found that the Δ indel regulates *IVD* expression in LCLs concordant with our luciferase results. Cell populations homozygous for the derived 5-bp deletion express *IVD* at significantly higher levels than those homozygous for the insertion, regardless of haplotype background (Fig. 3). This important step validates that the Δ indel modulates *IVD* expression endogenously, rather than merely having the capacity to regulate reporter expression on a vector backbone.

**Figure 3.**
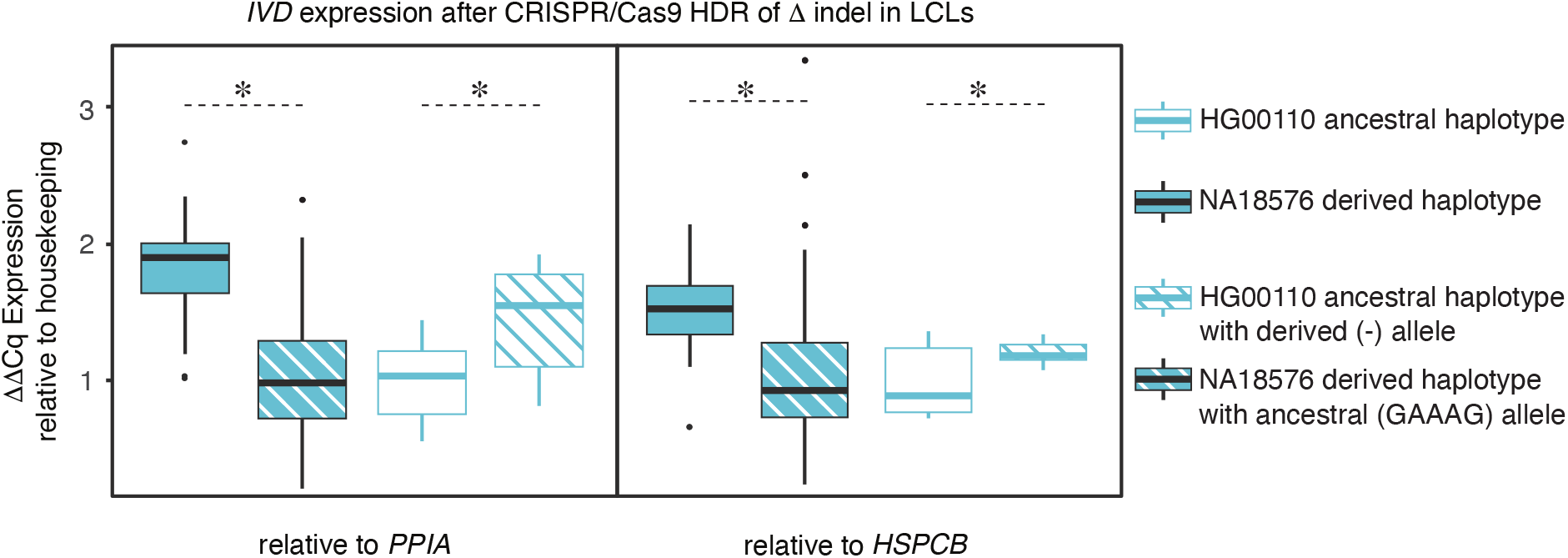
Reciprocally converting the Δ indel endogenously recapitulates the effect. We conducted CRISPR/Cas9 homology directed repair (HDR) to test the impact of endogenous replacement of the 5bp indel Δ on *IVD* expression relative to housekeeping genes in LCLs. We converted NA18576 LCLs from homozygous derived to homozygous ancestral with three technical replicates of three RNA extractions performed for each biological clone of the derived (2 clones) and ancestral (9 clones) genotypes. We converted HG00110 LCLs from homozygous ancestral to homozygous derived with five technical replicates of one RNA extraction performed for each biological clone of the ancestral (3 clones) and derived (3 clones) genotypes. We used *PPIA* and *HSPCB* genes for internal standards. *indicates p-value < 0.01 for 2-sample t-test.

### MPRA

Since the Δ indel alone does not fully explain the *IVD* eQTL association peak (the τ peak variant maintains p-value = 2.51e-24 with *IVD* expression after conditioning on the regulatory Δ indel), we utilized MPRA to comprehensively test variants across the region in the *IVD* locus at once. MPRA is capable of testing hundreds to thousands of putative enhancer elements together by sequencing barcodes placed in the 3’ UTR of the reporter gene. Expressed barcodes, each uniquely tagging a specific test sequencing, are counted to measure enhancer activity across many putative elements at once. MPRA enables the testing of a much larger set of variants including all alleles tightly linked with the τ variant, as well as all those which maintain conditional association of p-value < 1e-6 after controlling for the τ variant (referred to here as “conditional eQTLs”). The current MPRA, however, is limited in the amount of genomic context surrounding the variant that can be tested. In total, we tested 50 variants, centered within 180 bp of their native genomic context using MPRA. (See Supplementary Tables 1 and 2 for *IVD* eQTL and metabolite associations. See Supplementary Notes on the lack of conditional associations in the GWAS results.)

We first assessed how the MPRA approach, with its smaller testable genomic context (Fig. 4a), compared to our luciferase assays on the original eight τ-linked peak eQTLs of interest. MPRA agrees with our luciferase results, with neither the peak τ variant, its proxies, nor the regions that encompass them showing any evidence for activity (Fig. 4b; see Supplementary Table 5 for complete MPRA results).

**Figure 4.**
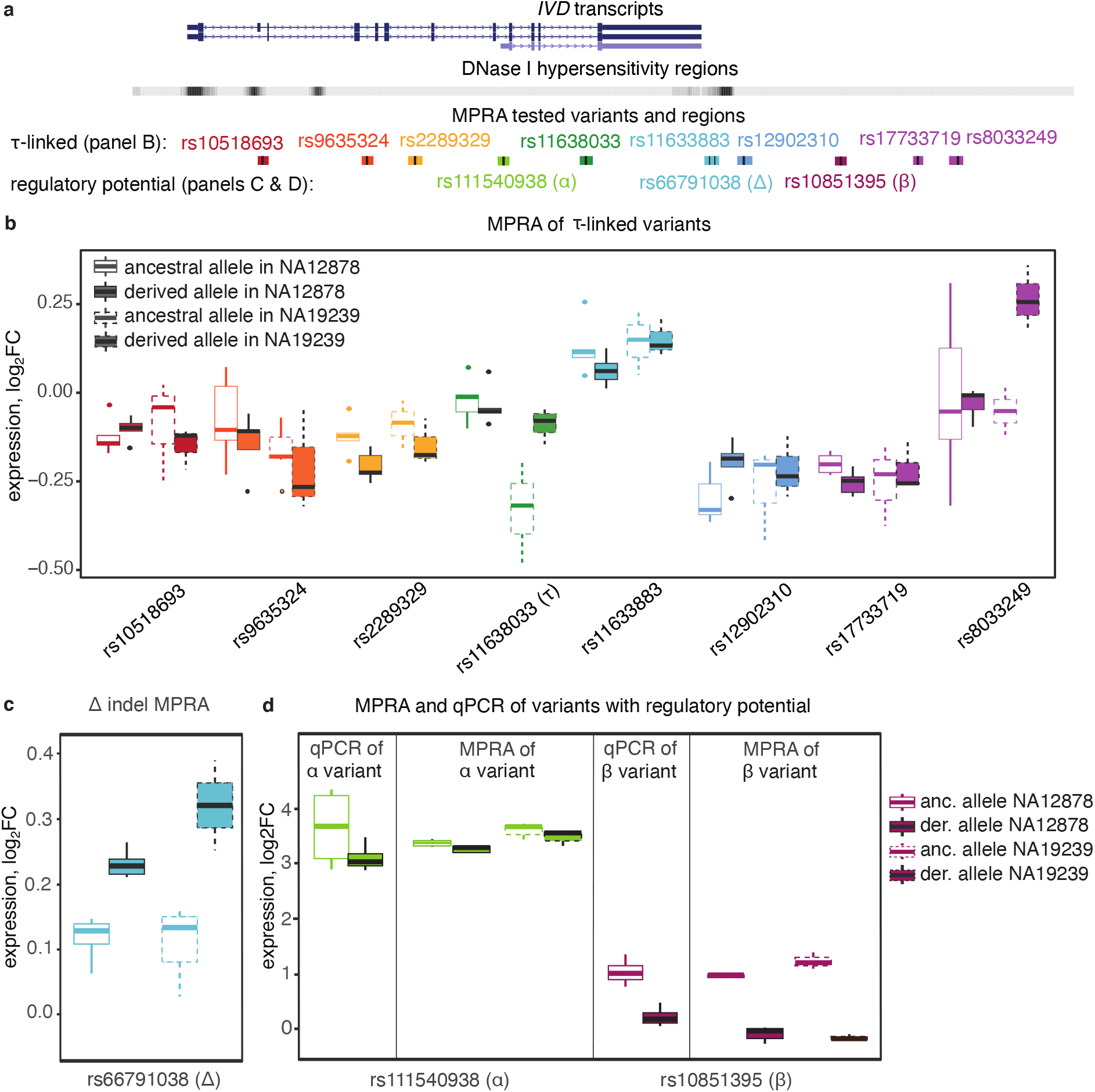
MPRA fails to identify any τ-linked variants, but identifies α and β variants with conditional association to *IVD*. **a** We show the positions with respect to the *IVD* transcripts of the same 8 SNPs linked with r^2^ > 0.9 to the peak associated *IVD* eQTL (referred to as τ-linked variants), along with the three variants that we detected to have regulatory potential by MPRA and single-plex qPCR of pGL4.23 constructs. We denote variants with color and show the 150bp of genomic context included in the MPRA assay for each. Note that the τ-linked variant rs11633883 is just 30bp from the rs66791338 Δ indel, so we test them in the same region in the MPRA using all combinations of ancestral and derived alleles, centering on each of the variants. **b** We conducted our MPRA experiment in both NA12878 and NA19239 LCLs. We plot expression as log_2_ of fold-change (log_2_FC) of the MPRA reporter gene count relative to the plasmid DNA count for ancestral and derived alleles of the τ-linked peak eQTLs in their respective MPRA constructs using boxplots. They have negligible enhancer activity (proximity to 0) and skew (comparing derived to ancestral) in both NA12878 and NA19239 LCLs. **c** We plot MPRA reporter expression (log_2_FC) of the Δ indel ancestral and derived constructs to show potential enhancer activity and allelic skew (derived over ancestral) of this variant. **d** We show the strong enhancer activity of the α region (both alleles) and the β variant (ancestral allele only) in MPRA using a boxplot of MPRA reporter expression relative to the plasmid DNA (log_2_FC). We also show this activity replicates in a single-plex assay qPCR assay of the same MPRA regions cloned into the pGL4.23 vector, plotting the log_2_FC of luciferase driven by the ancestral and derived variants relative to luciferase from a minimal promoter (with no enhancer region inserted) in NA12878 cells, 6 technical replicates each. For the MPRA results, the derived allele of the α variant is decreased relative to the ancestral allele (for the two LCL lines, combined log_2_FC = 3.49 and 3.37, −log_10_pvalue = 319.68 and 320.74 for ancestral and derived alleles, respectively; log_2_skew = −0.12, −log_10_pvalue = 6.42). The ancestral allele of the β variant is increased relative to the derived allele (combined log_2_FC = 1.13, −log_10_pvalue = 220.38 for the ancestral allele; combined log_2_FC = −0.07, −log_10_pvalue = 0.73 for the derived allele; log_2_skew = 1.20, −log_10_pvalue = 5.51).

We next examined the Δ indel in both orientations and on alternative haplotype backgrounds. We found it ranks third in our MPRA for allelic skew and replicates the direction of effect seen by luciferase (log skew = 0.15, p-value = 5.75e-4; Fig. 4c). However, as we expected given the lack of larger genomic context, the total activity of the Δ indel with its surrounding 180 bp is weaker, and it does not pass our threshold of significance (FDR = 0.106). To validate the impact of genomic context on enhancer activity of the Δ indel, we created seven pGL4.23 constructs containing the Δ indel with varying amounts of genomic context from the original 2.2kb insertion to the 150bp region used in the MPRA. As expected, the reduction of genomic context dramatically reduces the luciferase activity that we detected (Supplementary Fig. 2).

Applying MPRA to the other conditional eQTLs, we identify two variants with the potential for regulatory activity, rs111540938 (“α”) and rs10851395 (“β”) (Fig. 4a and d). These both give stronger signals of allelic skew than Δ indel, and are significant in our analysis. Specifically, both alleles of the α variant region drive overexpression of the reporter gene relative to the count of plasmid DNA for those constructs in the same cells, and the ancestral α allele having significantly increased expression compared to the derived. For the β variant, the ancestral allele shows increased expression relative to plasmid DNA count, while the derived allele has no activity by MPRA.

We validated our MPRA results by cloning α and β variant regions identical to those tested by MPRA, into a luciferase vector, and testing the regulatory activity of those constructs in NA12878 LCLs. qPCR of the luciferase gene from these single-plex assays clearly replicates the enhancer activity observed in the MPRA for these variants when we plot the log2FC of luciferase driven by the regulatory regions relative to luciferase driven by a minimal promoter alone, tested alongside in the same experiment (Fig. 4d). Interestingly, measuring luciferase activity, rather than gene expression directly by qPCR, of these three variants (α, β, and Δ) clearly replicates results only for the Δ indel and not the other two (Supplementary Fig. 3 and Fig. 2). Discrepancies in enhancer detection in MPRA and luciferase assays is an important area for future research. For example, both α and β are located within short interspersed nuclear elements (SINEs), which are common sites for transcriptional regulation^36^. The entire SINE surrounding each variant could not be included in the MPRA vector due to the size limitations. So perhaps the incomplete SINE element impacted where transcription started, and thus translation of luciferase. This also further highlights genomic context, and diverse potential signatures of regulatory importance in assay design for testing regulatory elements.

We then looked more closely at the evidence that the α and β could be driving *IVD* phenotype associations. We found that the β variant is the top associated variant in the region with bioavailable testosterone in a GWAS of European women^25^ and this SNP’s strongest eQTL association in GTEX is with *IVD* expression in skeletal muscle (p =1.2e-23, ancestral allele associated with increased expression). The α variant is not strongly linked to any GWAS peaks. At the same time, we note that rs2034650, the GWAS for pulmonary fibrosis^16^, and rs2304645, the GWAS for pulmonary function^18^, were among the conditional variants that we tested. They showed no activity by MPRA, but they are each tightly linked (r^2^ = 0.978 in GBR) to both the Δ indel and the β variant suggesting these might be the causal variants for the pulmonary associations.

### MPRA bashing

Since we had already identified the Δ indel and validated it in an endogenous context using CRISPR/Cas9 HDR, we also used our MPRA to try to localize the active nucleotides surrounding the regulatory indel using a bashing strategy. This approach has been used successfully with other loci to identify enhancer motifs^33,35^, but we recognized that lack of genomic context in MPRA may limit its efficacy for this particular region. In short, we created 1-bp and 5-bp deletions and single nucleotide mutations across the Δ indel region to test which local nucleotides are critical to drive expression of the MPRA reporter. We localized enhancer activity to the 15 nucleotides upstream of the Δ indel based on detecting lower expression when these sites were deleted on oligos containing the Δ derived deletion allele (Supplementary Fig. 4). This is consistent with our previous results that found the derived deletion drives increased expression over the ancestral allele. Unfortunately, as expected due to limited genomic context, the signal to noise of the assay was too low to recover the specific binding motif through saturation mutagenesis across the oligos. (See Supplementary Notes and Supplementary Fig. 2.)

### Functional haplotype analysis

After detecting three functional variants (α, β, and Δ), we wanted to understand how these variants fall on regional haplotypes, and whether they could potentially explain the isovalerylcarnitine GWAS and *IVD* eQTL peak variant (τ). All three functional variants retain association with *IVD* expression after controlling for τ, as shown when the conditional *IVD* eQTLs are plotted against *IVD* eQTLs (Supplementary Fig. 5). The higher expressing ancestral β and derived Δ alleles are highly correlated (r^2^ = 0.91) with each other in the Great Britain (GBR) population of 1000G phase 3 and occur at about 50% frequency (Fig. 5a). The α variant is tightly linked but weakly correlated with β and Δ (D’= 1 for each, r^2^ = 0.1 and 0.105 in GBR). The α derived allele which decreases expression, however, is always observed on the background of the β and Δ haplotype at 10% frequency in GBR.

**Figure 5.**
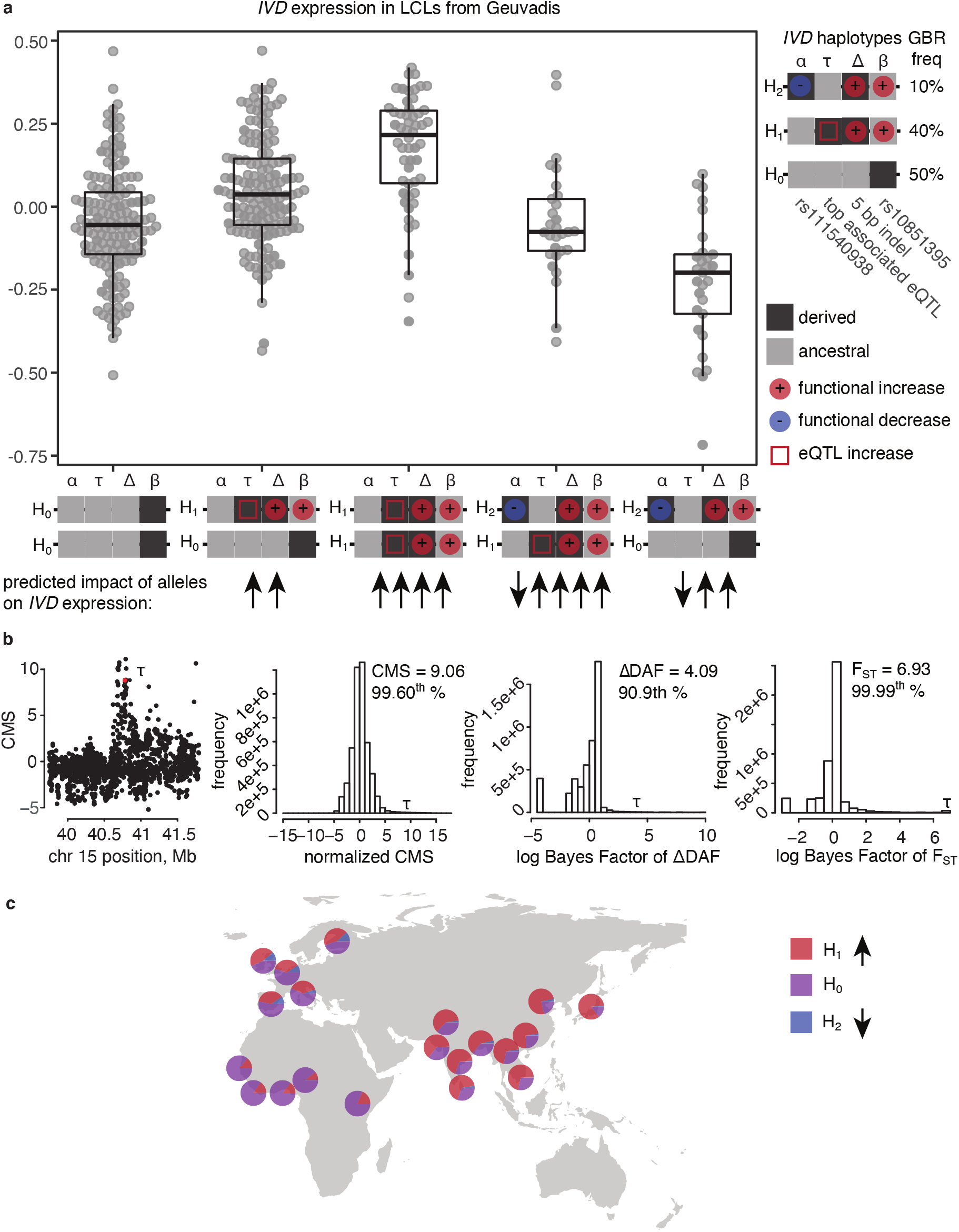
*IVD* functional variants generate a synthetic association of the haplotype tagged by the τ variant, which is positively selected in JPT. **a** We plot normalized *IVD* expression data from the Geuvadis RNA-seq study in LCLs using boxplots, dividing individuals by haplotypes of the potential regulatory variants, Δ, β, and α, that we detected. We find that these functional alleles predict *IVD* expression on haplotypes, tagged by the τ variant, across individuals. **b** We show that the τ variant, which tags the functional haplotype generated by *IVD*’s three regulatory variants, falls high in the genomewide distribution of SNPs for CMS component and composite scores in the JPT population. **c** We show world population frequencies of *IVD* functional alleles. This shows that *IVD* high expression alleles are at highest frequency in Japanese from phase 3 1000G.

The additive combination of these three variants regulating *IVD* could explain why the 40% of individuals with the Δ deletion, β ancestral, and α ancestral alleles have the highest expression of IVD in LCLs from European individuals in the Geuvadis RNA-seq study^14^ (Fig. 5a). Notably, this haplotype is highly correlated (R^2^ = 0.98) with the derived allele of τ, the peak eQTL and GWAS association. We moreover found no evidence that the τ variant nor its proxies have any regulatory potential by MPRA or luciferase assays. The data suggest that the combination of the three detected functional variants on population haplotypes generates a synthetic peak of association that drives the primary *IVD* eQTL and isovalerylcarnitine GWAS signal. As supporting evidence of this hypothesis, we note that after controlling for the τ variant, the associated effect sizes of the three functional variants change in concordance with our expectations based on functional evidence (Supplementary Table 2). However, we cannot exclude the possibility that other linked variants in the region also regulate *IVD*, but remain undetected by our functional assays.

We also analyzed the eQTL signals in GTEx (a database of eQTLs in many different tissue types) to see if they are consistent with the hypothesis that the regulatory activity of the α, β, and Δ variants generate a synthetic peak around the τ variant. We observed a pattern of association across tissues consistent with our hypothesis (Supplementary Fig. 6). Specifically, the Δ derived deletion and β ancestral alleles have the strongest positive associations with *IVD* in skeletal muscle (p = 1.2e-23), whereas the α derived allele has negative associations with *IVD* in most other tissues, excluding skeletal muscle. The τ derived allele, similar to β ancestral and Δ derived alleles, has a positive *IVD* association in skeletal muscle. The τ derived allele also has positive *IVD* associations in many of the tissues where the α derived allele has negative associations. (Note that the τ derived allele and the α derived allele are anti-correlated on population haplotypes, refer to Fig. 5a.) Therefore, τ’s association with *IVD* expression across tissue types is consistent with being the sum of the activity of our three regulatory variants (α, β, and Δ) acting with differing strengths across tissues.

We found a number of other associations in the GTEx data that highlight the pleiotropic nature of regulatory variants. The β variant also exhibits weaker associations to nearby *KNSTRN, DISP2*, and *BAHD1* genes in a variety of muscle, tibial, esophageal, and adipose tissues in GTEX. It further appears to be a splicing QTL for IVD in thyroid and fibroblasts in GTEX. Likewise, the top eQTL for *IVD*, the peak τ variant, is also an eQTL for *DISP2* and *KNSTRN* in a variety of tissues, and shows splicing QTL association for *IVD* in thyroid and fibroblasts. The α variant, furthermore, shows strong eQTL associations with *KNSTRN* in opposite direction of *IVD* in most of the tissues tested in GTEX. These pleiotropic effects make the connection between local regulatory variation and human phenotypes even more challenging.

### Dating and selection analysis for functional haplotypes and alleles

Finally, we revisited the signatures of positive selection in the Japanese population, closely examining the haplotype with increased *IVD* expression. We found that the peak eQTL variant that tags the high-expression haplotype (R^2^ = 0.98) is at highest frequency globally in the JPT, and has evidence of positive selection in the population. CMS places this SNP in the top 0.4% of SNPs with evidence for selection in JPT (Fig. 5b and Supplementary Table 3). This was largely driven by very high allele frequency differentiation, which is consistent with the actions of selection on a population (F_ST_ in the top 0.01% for JPT). In addition, perfect proxies for the Δ indel and β variant also have high allele frequency differentiation (F_ST_ in the top 1% for both variants) and fall in the top 1% for the overall CMS selection statistic in the JPT population.

Measures of selection specific to newly derived variants, such as those that detect long haplotype, are not high for our variants, fitting our expectation that these variants were not newly derived in the JPT population. Instead we estimate they arose prior to the human migration out of Africa (79,000 to 166,000 years old for Δ, and 291,000 to 432,000 years old for β). Overall, the JPT sample had higher frequency of alleles (Δ deletion, β ancestral, and α ancestral) conferring increased *IVD* expression and activity (Fig. 5c). Furthermore, the *IVD*-associated haplotype generated by these three regulatory alleles exhibits signatures consistent with positive selection on standing regulatory variation in JPT.

## DISCUSSION

By applying multiple complementary approaches, we detected three causal eQTLs, an indel and two SNPs, which appear to collectively regulate IVD expression, and have potential consequences for human metabolism and health. These regulatory variants map onto the GWAS signal for leucine metabolites in blood. While extreme perturbation to *IVD* causes fatal acidemia, we find positive selection in JPT increased the frequency of the haplotype with highest *IVD* expression. This aligns with other research showing repeated selection in diverse populations on variants regulating the lactase enzyme^37–39^. Many similar cases are likely to exist as selection signals are enriched among likely regulatory variants and metabolic genes as well^6,40^.

Our research suggests probing the effect of variable *IVD* metabolite ratios on human physiology across diverse populations, particularly in the tissues where the selected regulatory alleles appear to be active, would help us understand phenotypic relationships and consequences for population health. For example, the β functional variant is the peak of an association with testosterone levels in a GWAS of European women. The high-expression ancestral β allele, which is at highest frequency in East Asians, is associated with increased testosterone and increased *IVD* in skeletal muscle. Also, two of *IVD*’s regulatory variants, the Δ indel and the β variant, are closely linked to a nearby GWAS variant associated with pulmonary fibrosis in Europeans^16^ with the risk alleles at 84% in JPT. The relationship between testosterone levels, the oxidative damage or inflammatory pathway of pulmonary fibrosis, and leucine-related metabolism, signaling, and acidemia merit further study. We cannot exclude the possibility that selection could have been driven by pleiotropic impacts on other nearby genes associated with the same set of variants, such as *KNSTRN* (opposite direction of regulation) or *DISP2* (same direction as *IVD*). However, the IVD metabolite association suggests a reasonable phenotypic consequence of *IVD* regulation that could plausibly contribute to the observed selection signal and risk for disease.

The three linked regulatory alleles we detected also act in opposing directions and provides a compelling example where current models to decipher causal eQTLs *in silico* are unable to capture situations with complex genetic architecture. The linkage pattern of these alleles in the GBR population could explain why a haplotype tagged by the τ variant forms the peak of association for the eQTL and GWAS. While none of the peak-associated variants appear to be functional by MPRA and luciferase in our study, they are linked to sub-haplotypes that contain two positive functional alleles (Δ and β) and one negative functional allele (α). This could generate a synthetic association of τ with *IVD* expression and leucine metabolites. Our findings suggest population variation in leucine metabolite levels in whole blood is driven by the Δ and β variants regulating *IVD* expression in skeletal muscle, and by the α variant regulating *IVD* in multiple other tissues. The possibility remains that τ-linked variants could regulate *IVD* by a mechanism undetectable in episomal-based assays or in other tissues, creating an eQTL association shadow under which the detected functional alleles fall. However, the functional alleles we do detect by MPRA, luciferase, and CRISPR/Cas9 appear sufficient to explain the eQTL and GWAS peaks. Our findings should motivate future researchers to consider how multiple variants in linkage disequilibrium may control individual regulatory phenotypes. Our detection of the indel (Δ), in particular, also highlights the importance of probing less-studied types of variation, which may be nearby or linked to predicted candidates.

Our efforts to probe the regulatory landscape governing the gene *IVD* are part of our broader effort to develop a process for interrogating human non-coding genetic variation. Computationally, we analyzed the peaks of association with *IVD* and leucine metabolites for correlation, conditional association, and population-genetic signals of selection. Functionally, we applied luciferase assays of large regions of genomic context, CRISPR/Cas9 genome editing, and high-throughput MPRAs to resolve regulatory motifs and widely interrogate the region. We find that vector-based assays for detecting these regulatory variants are sensitive to the amount of genomic context and the vector backbone. Specifically, large regions of genomic context in the luciferase assays can amplify weaker signals from MPRA, but conversely MPRA applied broadly to eQTL peaks enables detection of buried regulatory signals. This highlights the importance of 1) combining multiple technologies to explore regulatory candidates and 2) further technical development to stabilize vector behavior, make assays able to accommodate more genomic context, and improve sensitivity of enhancer activity on different vector backbones. The research here demonstrates how large association studies, population-genetic statistics, and high-throughput, traditional, and emerging technologies can be combined to begin to parse complex regulatory loci critical to human health and fitness.

## METHODS

### Analyzing IVD eQTLs, isovalerylcarnitine association, and selection peak

The *IVD* eQTL peak was detected in the European population individuals in the Geuvadis RNA-seq study conducted in LCLs^14^. Data is available for download at https://www.ebi.ac.uk/arrayexpress/browse.html?keywords=E-GEUV*. The GWAS association peak surrounding variants in the *IVD* locus and isovalerylcarnitine, controlled for propionylcarnitine, was described in Suhre et al. 2011^41^ and Shin et al. 2014^15^. GWAS data is available for download at http://metabolomics.helmholtz-muenchen.de/gwas/. In order to analyze this association peak, we obtained genotyping and metabolite data from the Twins UK individuals used in these studies by application http://www.twinsuk.ac.uk/data-access/. We used the MERLIN program to conduct an association scan between the Twins UK individual genotypes, imputed to 1000G phase 3 SNPs, and isovalerylcarnitine normalized to propionylcarnitine. Genotyping and metabolite data from the Twins UK project were obtained by application to the Department of Twins Research at King’s College London. Genotyping for this project was provided for 5,654 individuals (2,040 on the HumanHap300, 3,461 on the HumanHap610Q, and 153 on the HumanHap1M and 1.2MDuo arrays). The Twins UK data management team imputed these genotyping data to 1000G phase 3 using the Michigan Imputation Server upon our request. Metabolite data for 5,004 individuals was also provided, allowing us to conduct the association on the subset of 4,678 individuals with both genotyping and metabolite data. We conducted the association analysis on variants in chromosome 15 within a 4.5 Mb window of the transcription start site of *IVD*, from 38,500,000 to 43,000,000 on chr15 in hg19, which encompasses the gene body and any potential regulatory elements upstream or downstream of the gene (Supplementary Table 1).

We examined genomic signatures of selection using the Composite of Multiple Signals (CMS)^6^ framework applied to 1000G phase 3 individuals from the JPT population. Briefly, the CMS statistic at a given SNP is the log-normalized product of a set of Bayes factors, which each indicate the relative probability of selection for a semi-independent input statistic (iHS, deliHH, nSL, H12, XP-EHH, F_ST_, and delDAF), as generated from simulated data. This updated application of CMS to phase 3 1000G individuals will be fully described in a forthcoming publication (Vitti, unpublished research). UCSC tracks for selection statistics from this new application in 1000G phase 3 populations are available in bigwig format at https://personal.broadinstitute.org/vitti/ucsc012918/.

We used linear regression in R to regress the metabolite association peak on the *IVD* eQTLs from Geuvadis (Supplementary Table 2), limiting our regression analysis to those variants in the peak of either association with p-value < 1e-6. Similarly we used linear regression in R to regress the composite selection score from CMS in the JPT phase 3 1000G (Supplementary Table 3) on the *IVD* eQTLs in LCLs from Geuvadis. For the conditional associations with *IVD* expression and isovalerylcarnitine (controlled for propionylcarnitine) the top associated variant for each was used as a covariate in the association analysis (Supplementary Tables 1 and 2).

### Luciferase reporter assays

We selected variants within r^2^ = 0.9 of the top metabolite association and eQTL in the Geuvadis dataset, and PCR amplified surrounding regions with ancestral/derived alleles of the eQTL variants from NA12812 genomic DNA, using DNase I, histone modifications, and conservation to define boundaries of the regions. The regions ranged in size from ~600-bp to 2-kb. We cloned these regions into the minimal promoter, luciferase reporter vector using SfiI restriction digestion and ligation to insert each region just upstream of the minimal promoter (primers provided in the Supplementary Table 4). We conducted 6 replicates of 1μg firefly vector for each construct into ~5 × 10^5^ NA12878 LCLs in 10μL Neon electroporation reactions (1200V, 20ms, 3 pulses), including 100ng renilla vector (pGL4.74) as an internal control. The NA12878 were plated in 100μL prewarmed RPMI + 15% FBS in a 96-well plate. We measured luciferase luminescence for firefly and renilla using the Dual Glo Luciferase kit 24 hours post transfection and the Spectramax L to read luminescence, in order to compare expression driven by ancestral eQTL constructs to derived.

We additionally tested the ancestral and derived haplotype vectors containing the 2.2-kb *IVD* downstream region showing a significant difference in NA12878 LCLs in HEK293 and HepG2 cells. In HEK293 cells, six replicates for each construct of 100ng firefly vector plus 10ng renilla vector was transfected into ~5 × 10^4^ cells, seeded 24 hours prior at 2.5 × 10^4^ cells/well in 100μL DMEM + 10% FBS and grown to 70% confluence, using Lipofectamine 2000 transfection reagents and DMEM without FBS. In HepG2 cells, six replicates for each construct of 100ng firefly vector plus 10ng renilla vector was transfected into ~5 × 10^4^ cells, seeded 24 hours prior at 2.5 × 10^4^ cells/well in 100μL MEM + 10% FBS and grown to 70% confluence, using Lipofectamine 3000 transfection reagents and MEM without FBS. Luciferase luminescence was measured using the Dual Glo Luciferase kit 24 hours post transfection as previously described.

Subsequent to our MPRA, primers were designed to amplify and clone oligos containing the rs111540938, rs10851395, and rs66791338 variants from our MPRA library into the luciferase vector backbone. The regions containing rs111540938 and rs10851395 were cloned using a SfiI restriction digest and ligation strategy as described above. The region containing rs66791338 was cloned using a Gibson assembly reaction into a linearized PCR product of, using a 15:1 molar excess of insert:backbone. Luciferase assays for these constructs was conducted in NA12878 LCLs, and luminescence was measured as described above. Gene expression was also measured after RNA extraction by RNeasy miniprep and Turbo DNase by qPCR using primers for firefly luciferase, controlled for renilla luciferase (all primers provided in the Supplementary Table 4).

### Site directed mutagenesis

Using primers complementary to the 10 variants differing between the tested constructs, we PCR amplified the vector containing the derived rs11633883 haplotype construct from NA12812, converting individual variants to the version on the ancestral rs11633883 vector construct. In these PCRs we annealed at 60C for 1 minute and extended at 72C for 1 minute/kb of construct for 18 cycles. We tested each of the 10 newly created vectors compared to the original ancestral and derived rs11633883 versions of the vector using the luciferase reporter assay described above in HEK293 cells. We also used site directed mutagenesis as above to convert the rs66791338 insertion ancestral allele on the starting ancestral rs11633883 vector to the derived deletion allele, and tested ancestral to derived and derived to ancestral constructs using the luciferase reporter assay in NA12878 LCLs.

We also used site-directed mutagenesis to delete portions of the vector with the 2.2-kb insert from the 3’ region downstream of *IVD* containing the rs11633883 and rs66791338 variants. We used site-directed mutagenesis, as well, to insert a 66-bp region from the MPRA vector backbone, residing between the minimal promoter and the oligo insertion site, into the luciferase vector at the analogous position, causing the vectors to be identical with the exception of the luciferase versus GFP open-reading frame (all primers provided in the Supplementary Table 4).

### CRISPR/Cas9 homology directed replacement of rs66791338

For both HG00110 LCLs homozygous ancestral and NA18576 LCLs homozygous derived for rs66791338, we transfected ~5 × 10^5^ LCLs with Cas9 protein and tracrRNA/crRNA duplex (18.75 nmoles each) specific to the ancestral or derived version, along with 50 nmoles ssDNA oligo containing the replacement allele and 90-bp flanking arms homologous to the region. We conducted transfections in 10 μL Neon electroporations (1600 V, 10 ms, 3 pulses) and plated cells in 500 μL RPMI + 15% FBS in a 24-well plate and incubated at 37C. After 3 days DNA was harvested from half of the cells using Qiagen DNeasy columns, and the surrounding region was amplified and sequenced on a MiSeq to obtain NHEJ and HDR efficiency. Upon confirming NHEJ and HDR efficiency, LCLs were diluted to ~2 per 25μL of RPMI + 30% conditioned media + 15% FBS (filtered through 2μm flask to remove cellular contamination) and plated in 4-8 × 384-well plates. These were grown in isolation in an incubator for 1 week. 25μL of additional RPMI + 30% conditioned media + 15% FBS was added to each well and LCLs were grown for an additional 4-7 days. Wells with growth were then marked and transferred with 50μL additional RPMI + 15% FBS to a 96-well plate. After 1 day of growth 50μL additional media was added. After another day of growth, 75μL of LCLs + media were lysed for 1 hour at 95C in 2x lysis buffer. 1μL of lysate was added to a 10μL PCR rxn to amplify the surrounding region. We used TruSeq dual indexing to prepare a library of the clones and sequence them on a MiSeq to assess genotype. Unmodified and HDR homozygous LCLs were expanded and confirmed by Sanger traces. RNA was extracted using MagMax RNA extraction, converted to cDNA using Superscript III, and expression of IVD was assessed relative to the housekeeping genes, *PPIA* and *HSPCB* (primers and guides provided in the Supplementary Table 4).

### MPRA strategy

We selected variants within r^2^ = 0.9 in the EUR population of 1000G phase 3 of top the top GWAS and eQTL signal, rs11638033, to include in our MPRA screen. We selected conditional hits to test by taking variants within a 50kb window surrounding the transcription start site of *IVD* with p-value < 1e-6 after conditioning on rs11638033. Our MPRA protocol is described in Tewhey et al. 2016^34^. We synthesized oligos using Agilent Technologies, which centered 180-bp regions around each allele of each variant, with an additional 15-bp of adapter on either side of the regions. We added 20-bp unique adapter sequences to the oligos using emulsion PCR, along with a constant region used for Gibson assembly into a linearized vector backbone. After amplifying the library in *E. coli*, we used 2 × 150-bp HiSeq reads to associate oligos with adapter sequence. Then, we obtained our final MPRA library by restriction digesting the library and using Gibson assembly to insert a minimal promoter and the GFP gene between the variant regions and the unique adapters so that the adapters land in the 3’ region of the GFP transcript. This library we electroporated into NA12878 LCLs using the Neon system (1200V, 20ms, 3 pulses) by Life Technologies in five separate replicates on sequential days. We also performed three replications in NA19239 LCLs. Each replicate, consisting of ~5 × 10^8^ LCLs was spun down, washed, and frozen at −80C 24 hours after transfection. We lysed LCLs by passing them through the needle of a syringe and harvested RNA from each replicate using RNA midiprep and digesting DNA using Turbo DNase, capturing GFP using DNA probes, and synthesizing cDNA. Finally, we added Truseq Illumina adapters to the cDNA, along with our starting MPRA library, and sequenced the unique adapters in a 1 × 30-bp HiSeq run (all primers provided in the Supplementary Table 4). We applied Benjamini-Hochberg to adjust for multiple testing. The threshold for significance in this assay was FDR < 0.05.

### MPRA bashing strategy of rs66791338 indel

We designed oligo pools containing single base pair and 5-bp deletions in a sliding window on the oligo containing rs66791338 ancestral and derived alleles, beginning 40-bp upstream of the variant and ending 25-bp downstream of the variant. MPRA libraries were constructed as described above. 4 × 500ng of library was electroporated into ~2 × 10^7^ NA12878 LCLs in four reactions according to the described conditions, which were later combined. Five replicate transfections of the library were performed on sequential days. RNA was harvested and analyzed according to the MPRA protocol described (primers provided in the Supplementary Table 4).

### Functional haplotype analysis

We analyzed the structure of haplotypes to see where the three functional SNPs lie with respect to SNPs tagging the GWAS and eQTL signal in the GBR population from 1000G phase 3 individuals. We sorted haplotypes and identified tagging variants using K-means clustering in R for the local region on chromosome 15: 40650207-40728146 in the hg19 build of the human genome using phase 3 1000G variants in the GBR population. We also sorted and plotted European Geuvadis expression data from LCLs based upon phased allelic combinations in individuals. Finally we plotted the frequencies of identified functional haplotypes in phase 3 1000G populations on a world map using Ben Fry’s Diaspora Program.

### Dating and selection analysis for functional alleles

We compared test statistics of selection from the Composite of Multiple Signals (CMS) test^6^ applied to 1000G phase 3 individuals for variants tagging the active haplotype and the functional regulatory variants to the genome-wide distribution as calculated for the JPT population (Vitti, unpublished research).

In order to estimate the ages of derived functional alleles, we used the haplotype-based method described in Stephens et al. 1998^42^ applied to the West African populations from 1000G phase 3, since the alleles do not appear to be on a selected haplotype in these populations. First, haplotype length was visualized for these populations by observing the chunk of alleles with no recombination surrounding the derived functional variants. Then, the fraction of haplotypes lost with the addition of more SNPs on each side of the derived variants were averaged on each side of the allele. This generated an estimate of haplotype age on either side of the functional derived alleles using a 25-year generation time. Since the rs111540938 derived allele was only present in one Gambian individual in the 1000G phase 3 dataset, we applied the same test in the European populations, where the derived T allele is at its highest frequency of 10%.

## DATA AVAILABILITY

UCSC genome browser tracks for the CMS selection statistics for the unpublished analysis of phase 3 1000G data are available in bigwig format at https://personal.broadinstitute.org/vitti/ucsc_012918/.

## ACKNOWLEDGEMENTS

We acknowledge funding from the Howard Hughes Medical Institute and the NIH grants DP2OD006514 to P. C. S. and K99HG008179 to R. S. T. The Cora Du Bois fellowship supported E. A. B. We thank Terence Capellini and members of the Sabeti lab for helpful guidance during the execution of this project.

## AUTHOR CONTRIBUTIONS

E. A. B., R. S. T., and P. C. S. conceived and designed this project. E. A. B., S. K., M. J. B., and R. S. T. performed experiments for this work. E. A. B., S. K., M. J. B., J. V., D. K., S. F. S., and R. S. T. analyzed data for this project. E. A. B., R. S. T., S. F. S., and P. C. S. were involved in the writing of this work. R. S. T. and P. C. S. co-supervised this work. P. C. S. is the corresponding author for this work.

## ETHICS DECLARATIONS

We have no conflicts of interest to declare.

## Notes

https://personal.broadinstitute.org/vitti/ucsc_012918/

### Competing Interest Statement

The authors have declared no competing interest.

## REFERENCES

1. 1000 Genomes Project Consortium et al. A global reference for human genetic variation. Nature 526, 68–74 (2015).

2. Lek, M. et al. Analysis of protein-coding genetic variation in 60,706 humans. Nature 536, 285–291 (2016).

3. Altshuler, D., Daly, M. J. & Lander, E. S. Genetic mapping in human disease. Science 322, 881–888 (2008).

4. Ward, L. D. & Kellis, M. Interpreting noncoding genetic variation in complex traits and human disease. Nat. Biotechnol. 30, 1095–1106 (2012).

5. Maurano, M. T. et al. Systematic localization of common disease-associated variation in regulatory DNA. Science 337, 1190–1195 (2012).

6. Grossman, S. R. et al. Identifying recent adaptations in large-scale genomic data. Cell 152, 703–713 (2013).

7. Dimas, A. S. et al. Common regulatory variation impacts gene expression in a cell type-dependent manner. Science 325, 1246–1250 (2009).

8. Corradin, O. et al. Combinatorial effects of multiple enhancer variants in linkage disequilibrium dictate levels of gene expression to confer susceptibility to common traits. Genome Res. 24, 1–13 (2014).

9. Tanaka, K., Budd, M. A., Efron, M. L. & Isselbacher, K. J. Isovaleric acidemia: a new genetic defect of leucine metabolism. Proc. Natl. Acad. Sci. U. S. A. 56, 236–242 (1966).

10. Sun, X. & Zemel, M. B. Leucine and calcium regulate fat metabolism and energy partitioning in murine adipocytes and muscle cells. Lipids 42, 297–305 (2007).

11. Li, F., Yin, Y., Tan, B., Kong, X. & Wu, G. Leucine nutrition in animals and humans: mTOR signaling and beyond. Amino Acids vol. 41 1185–1193 Preprint at https://doi.org/10.1007/s00726-011-0983-2 (2011).

12. Rachdi, L., Aïello, V., Duvillié, B. & Scharfmann, R. L-leucine alters pancreatic β-cell differentiation and function via the mTor signaling pathway. Diabetes 61, 409–417 (2012).

13. Grossman, S. R. et al. A composite of multiple signals distinguishes causal variants in regions of positive selection. Science 327, 883–886 (2010).

14. Lappalainen, T. et al. Transcriptome and genome sequencing uncovers functional variation in humans. Nature 501, 506–511 (2013).

15. Shin, S.-Y. et al. An atlas of genetic influences on human blood metabolites. Nat. Genet. 46, 543–550 (2014).

16. Fingerlin, T. E. et al. Genome-wide association study identifies multiple susceptibility loci for pulmonary fibrosis. Nat. Genet. 45, 613–620 (2013).

17. Allen, R. J. et al. Genome-Wide Association Study of Susceptibility to Idiopathic Pulmonary Fibrosis. Am. J. Respir. Crit. Care Med. 201, 564–574 (2020).

18. Shrine, N. et al. New genetic signals for lung function highlight pathways and chronic obstructive pulmonary disease associations across multiple ancestries. Nat. Genet. 51, 481–493 (2019).

19. Lam, M. et al. Large-Scale Cognitive GWAS Meta-Analysis Reveals Tissue-Specific Neural Expression and Potential Nootropic Drug Targets. Cell Rep. 21, 2597–2613 (2017).

20. Davies, G. et al. Study of 300,486 individuals identifies 148 independent genetic loci influencing general cognitive function. Nat. Commun. 9, 2098 (2018).

21. Hill, W. D. et al. A combined analysis of genetically correlated traits identifies 187 loci and a role for neurogenesis and myelination in intelligence. Mol. Psychiatry 24, 169–181 (2019).

22. Lee, J. J. et al. Gene discovery and polygenic prediction from a genome-wide association study of educational attainment in 1.1 million individuals. Nat. Genet. 50, 1112–1121 (2018).

23. de la Fuente, J., Davies, G., Grotzinger, A. D., Tucker-Drob, E. M. & Deary, I. J. A general dimension of genetic sharing across diverse cognitive traits inferred from molecular data. Nat Hum Behav 5, 49–58 (2021).

24. Demange, P. A. et al. Investigating the genetic architecture of noncognitive skills using GWAS-by-subtraction. Nat. Genet. 53, 35–44 (2021).

25. Ruth, K. S. et al. Using human genetics to understand the disease impacts of testosterone in men and women. Nat. Med. 26, 252–258 (2020).

26. Haas, C. B., Hsu, L., Lampe, J. W., Wernli, K. J. & Lindström, S. Cross-ancestry Genome-wide Association Studies of Sex Hormone Concentrations in Pre- and Postmenopausal Women. Endocrinology 163, (2022).

27. Tanaka, K. Isovaleric acidemia: personal history, clinical survey and study of the molecular basis. Prog. Clin. Biol. Res. 321, 273–290 (1990).

28. Dickinson, M. E. et al. High-throughput discovery of novel developmental phenotypes. Nature 537, 508–514 (2016).

29. Schlune, A., Riederer, A., Mayatepek, E. & Ensenauer, R. Aspects of Newborn Screening in Isovaleric Acidemia. Screening 4, 7 (2018).

30. Ow, D. W. et al. Transient and stable expression of the firefly luciferase gene in plant cells and transgenic plants. Science 234, 856–859 (1986).

31. Cong, L. et al. Multiplex genome engineering using CRISPR/Cas systems. Science 339, 819–823 (2013).

32. Mali, P. et al. RNA-guided human genome engineering via Cas9. Science 339, 823–826 (2013).

33. Melnikov, A. et al. Systematic dissection and optimization of inducible enhancers in human cells using a massively parallel reporter assay. Nat. Biotechnol. 30, 271–277 (2012).

34. Tewhey, R. et al. Direct Identification of Hundreds of Expression-Modulating Variants using a Multiplexed Reporter Assay. Cell 165, 1519–1529 (2016).

35. Patwardhan, R. P. et al. Massively parallel functional dissection of mammalian enhancers in vivo. Nat. Biotechnol. 30, 265–270 (2012).

36. Chuong, E. B., Elde, N. C. & Feschotte, C. Regulatory activities of transposable elements: from conflicts to benefits. Nat. Rev. Genet. 18, 71–86 (2017).

37. Myles, S. et al. Genetic evidence in support of a shared Eurasian-North African dairying origin. Hum. Genet. 117, 34–42 (2005).

38. Tishkoff, S. A. et al. Convergent adaptation of human lactase persistence in Africa and Europe. Nat. Genet. 39, 31–40 (2007).

39. Enattah, N. S. et al. Independent introduction of two lactase-persistence alleles into human populations reflects different history of adaptation to milk culture. Am. J. Hum. Genet. 82, 57–72 (2008).

40. Fraser, H. B. Gene expression drives local adaptation in humans. Genome Res. 23, 1089–1096 (2013).

41. Suhre, K. et al. Human metabolic individuality in biomedical and pharmaceutical research. Nature 477, 54–60 (2011).

42. Stephens, J. C. et al. Dating the Origin of the CCR5-Δ32 AIDS-Resistance Allele by the Coalescence of Haplotypes. Am. J. Hum. Genet. 62, 1507–1515 (1998).

