## Supplementary Figures 1-6 and Notes for "Three linked opposing regulatory variants under selection associate with *IVD*"

### SUPPLEMENTARY NOTES

#### No conditional associations in the GWAS results

We note that there were no additional variants from the metabolite GWAS peak to test using MPRA. The corresponding conditional association variants from *IVD*'s eQTL peak fail to maintain association with isovalerylcarnitine after controlling for the peak variant in the GWAS. We believe this occurs because a key haplotype in the region at 10% frequency in EUR was not tagged by any SNP in the low coverage genotyping array used for most of the Twins UK individuals. Therefore, we had no power to tease apart hidden independent associations among linked variants in the GWAS dataset.

#### High noise to signal of the MPRA bashing experiment

While we detected lower expression driven by oligos with deletions near the Δ derived deletion allele, the signal to noise of the assay was too low to recover the binding motif through saturation mutagenesis across the oligos, though this has been successful for other loci. Testing further, we found the magnitude of expression driven by the Δ deletion depends upon both the genomic context included – perhaps owing to a CTCF ChIP-seq peak nearby – and the vector backbone used in the MPRA versus luciferase assays (**Supplementary Figure 2**). An important technical avenue for future research should be to study how sensitive MPRA is to the amount of genomic context and the vector backbone.

**Supplementary Figure 1: Process for parsing important regulatory regions in the genome.**

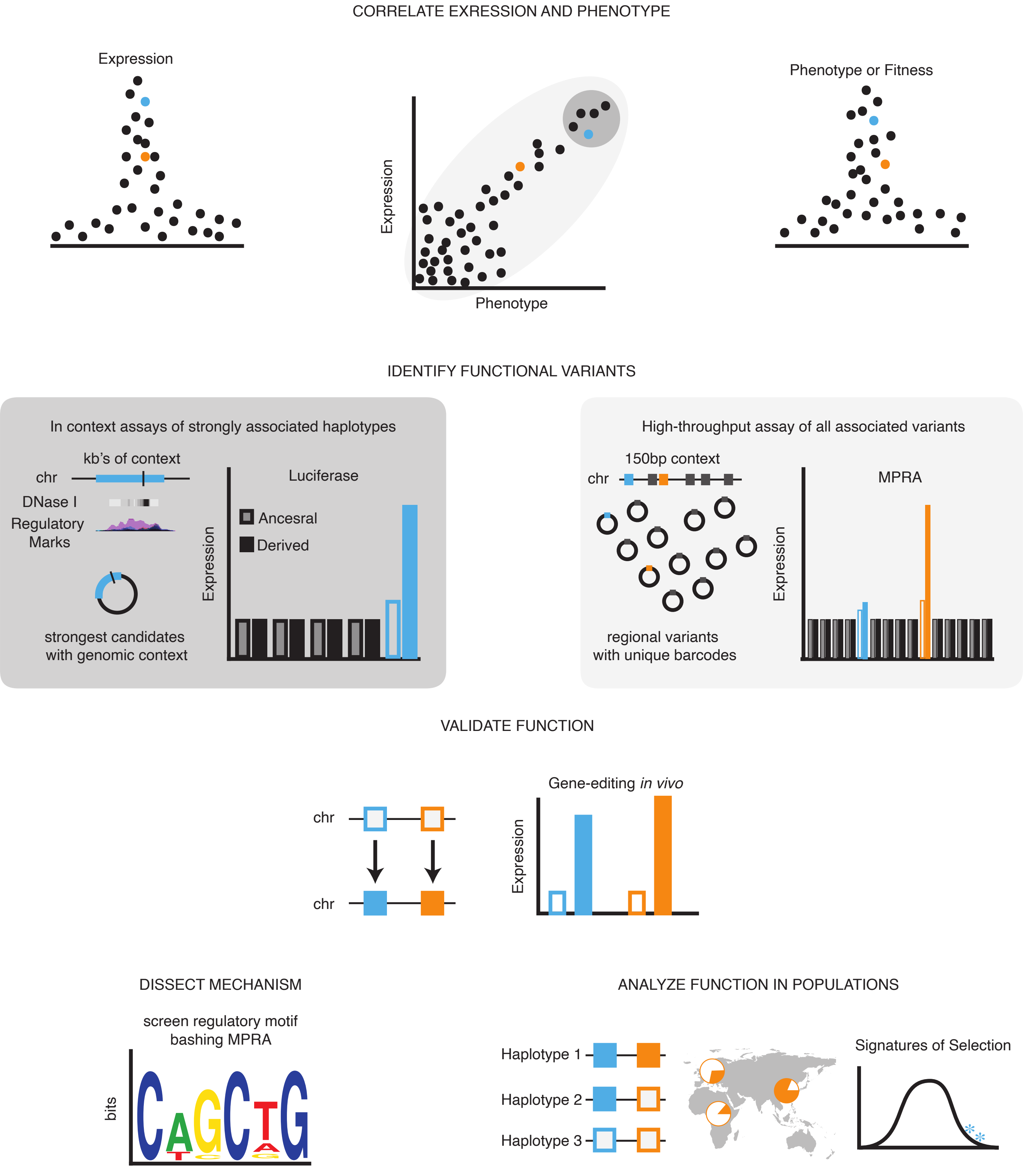

**Supplementary Figure 2: Amount of genomic context and vector backbone affect enhancer activity of the Δ deletion allele measured by luciferase**

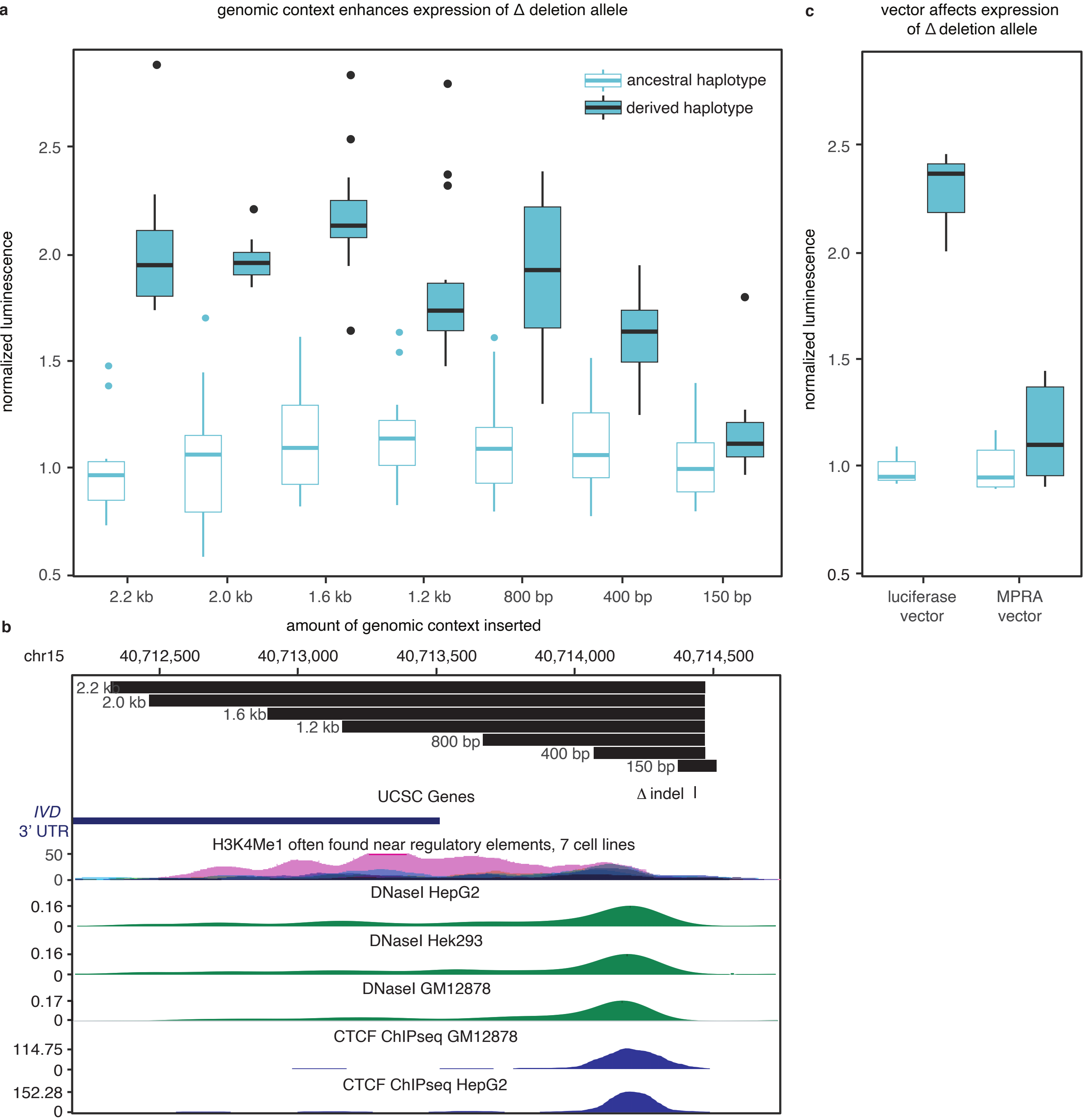


**a** We plot normalized luminescence (firefly luciferase to renilla luciferase) of pGL4.23 constructs containing the Δ indel region with genomic context surrounding ancestral insertion and derived deletion ranging from 150bp (identical to the MPRA insertion) to 2.2kb (original insert size). We show the effect size of the Δ deletion allele depends on having more genomic context since expression drops off rapidly for constructs containing 150bp of context compared to all larger amounts. **b** We show the construct insert sizes relative to the 3' of the IVD construct and regulatory marks, including H3K4Me1, DNase I hypersensitivity, and CTCF Chip-Seq from ENCODE. These regulatory marks are consistent with the larger regions containing important binding sites for enhancer activity. **c** We use boxplots of normalized luminescence of the 2.2kb region inserted into the pGL4.23 backbone and the MPRA vector backbone. High expression of the deletion allele only occurs from the pGL4.23 backbone. The MPRA vector contains 66-bp of additional sequence between the test sequence and the transcription start site (used for cloning test sequences into the vector backbone), which may impact the transcription start site.

### Supplementary Figure 3: Luciferase and MPRA match for the Δ indel, but not for the α and β variants.

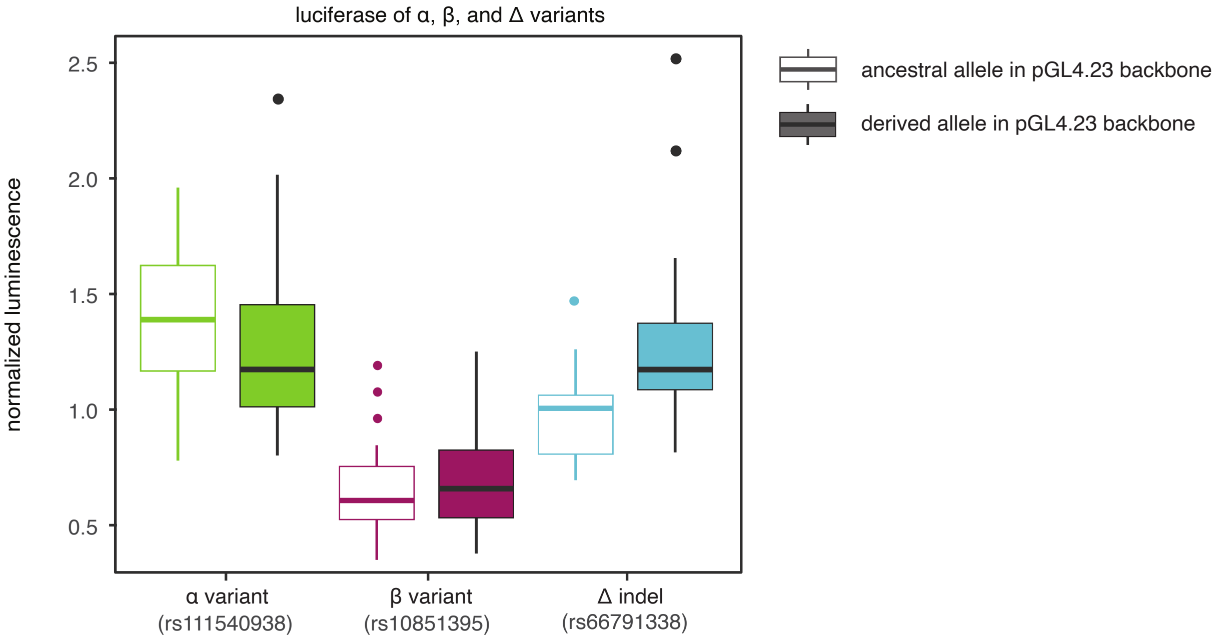


We use boxplots to show normalized luminescence (firefly luciferase normalized to renilla luciferase) of pGL4.23 constructs containing the MPRA inserts surrounding the α, β, and Δ variants. Boxes are colored by variant, as in Figure 4 with ancestral alleles in transparent boxes and derived alleles in opaque boxes. These luminescence results only replicate the effects observed in the MPRA for the Δ indel. However, qPCR of the same constructs do replicate the MPRA results, as shown in Figure 4. The α variant region does show the correct direction of effect for the alleles (ancestral increased expression relative to derived), but the magnitude of enhancer activity of this region appears dramatically reduced when measured by normalized luminescence rather than qPCR of luciferase.

**Supplementary Figure 4: Deletions near the derived Δ indel deletion allele diminish expression in bashing MPRA.**

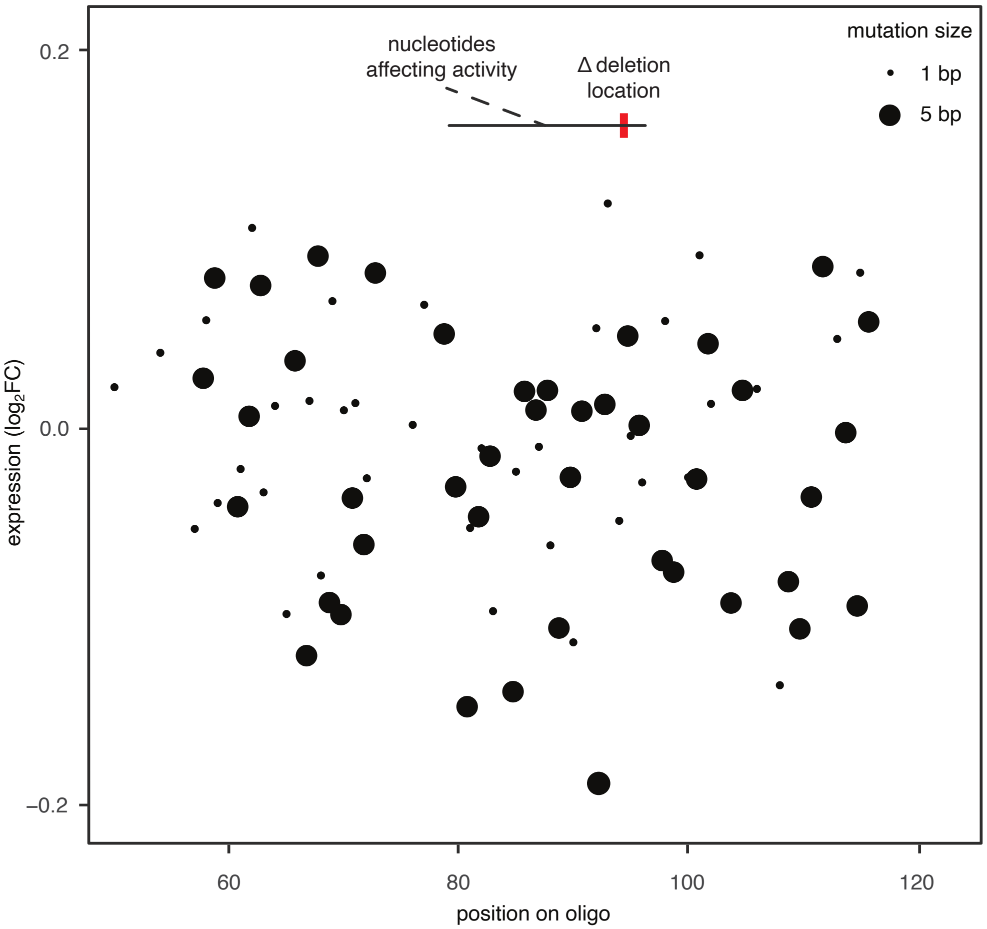


In our MPRA library we tested oligo inserts containing 1-bp and 5-bp deletions across the Δ indel region. We plot expression in log2FC on the y-axis for those deletion oligos (1-bp small dots; 5-bp large dots) relative to full-length oligos containing the Δ deletion allele. On the x-axis deletion oligos are plotted by position on the oligo with a red bar at top indicating the position of the Δ deletion location. Our results suggest a region just upstream of the Δ deletion location as important for enhancer activity, as deletions in this location produce a dip in expression on average relative to other deletions. However, the small effect size of the Δ indel in the MPRA makes us unable to dissect any transcription factor motif above the noise of the assay.

**Supplementary Figure 5: *IVD* functional variants maintain association with *IVD* expression after controling for τ, the top associated eQTL in LCLs**

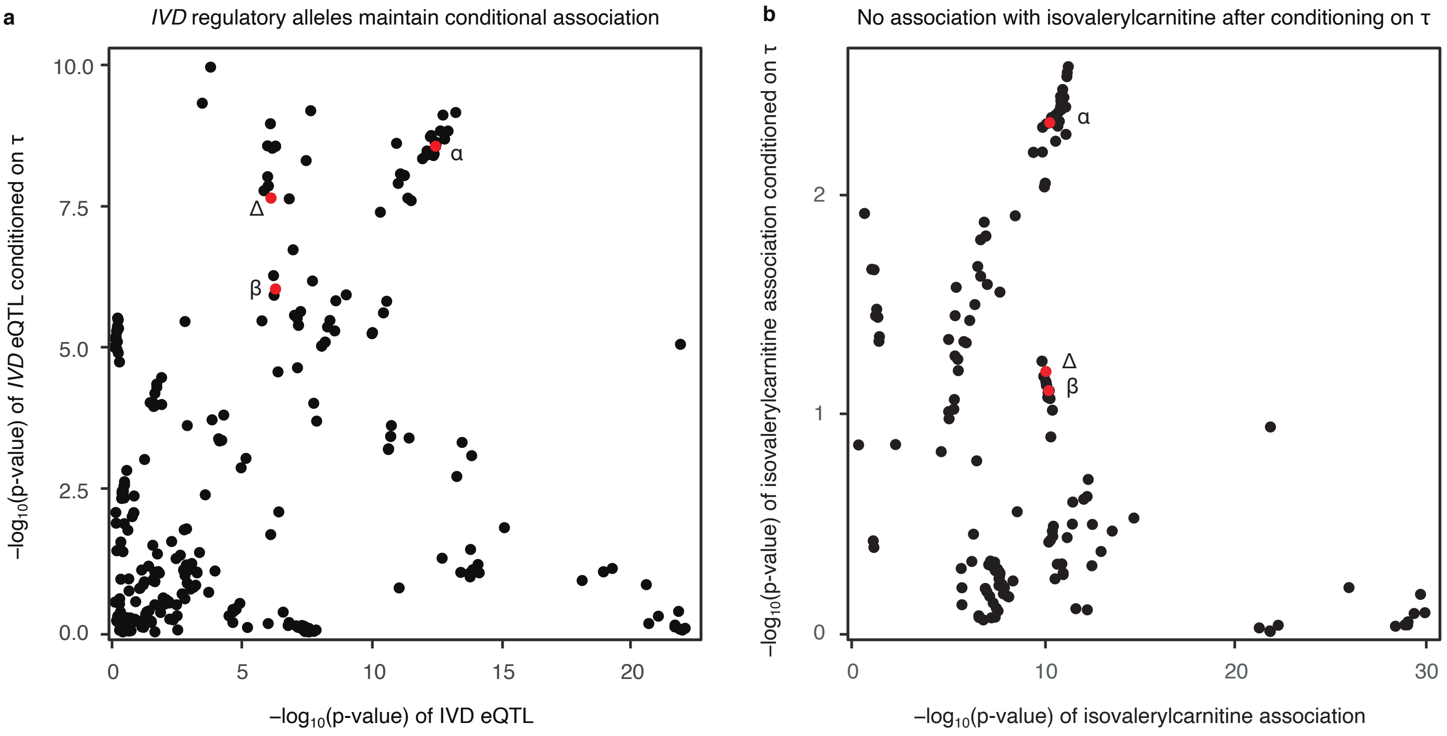


**a** The α, β, and Δ variants maintain conditional association with *IVD* expression after controlling for the τ peak eQLT variant. **b** Conditional association of these variants is not maintained in association with the isovalerylcarnitine metabolite. However, the genotyping panel used in this metabolite association did not include any proxies for the α variant. Therefore, we cannot detect conditional association of the three imputed functional variants in the metabolite association analysis.

**Supplementary Figure 6: The τ variant associates with expression of IVD in some of the same tissues as the α, β, and Δ functional variants**

α variant
τ variant
Δ indel
β variant

We mark variants of interest in the GTEx portal's locus browser for eQTLs of the IVD gene. Red dots indicate upregulation and blue dots indicate downregulation with color intensity indicating the Normalized Effect Size calculated in the database for a given tissue. Size of circle indicates the strength of the p-value (larger circles have lower p-values). We show that the α, β, and Δ functional variants associate with expression, along with the τ variant, which is the top eQTL in the Geuvadis eQTL study in LCLs. Specifically, the α variant associates in many tissues while the β and Δ variants seem specific to skeletal muscle in GTEX. The τ variant is associated both in skeletal muscle, where β and Δ appear to be active, and tissues where the α variant appears to be active
